# Discrete Ollivier-Ricci Flow Finds Distinct Subpopulations of Patients Treated with PD-1 Inhibition

**DOI:** 10.1101/2024.08.08.606714

**Authors:** James Bannon, Charles R. Cantor, Bud Mishra

## Abstract

In recent years immune checkpoint inhibitors (ICIs), also called immune checkpoint blockers, have revolutionized the standard of care for patients with cancers of many types. Researchers across many disciplines have endeavored to find biomarkers of response to ICI therapy but so far little consensus has been reached. In this paper we attempt to cluster patients in an unsupervised manner using discrete Ollivier-Ricci Flow (ORF). Our method surfaces populations with distinct survival curves which in turn allows us to find many potential biomarkers, including gene expression modules. We believe the algorithm may be of independent interest for clustering other datasets in a diverse set of research areas.

## 1 Introduction

In recent years approaches to treating cancer that leverage the immune system — called *immunotherapies* — have revolutionized the standard of care in many types of malignancies [1, 2, 3]. One class of immunotherapy, called immune checkpoint inhibitors (ICIs), has been successful in a diverse range of solid tumors [4]. Checkpoint inhibitors can be tremendously effective, leading to greater tumor shrinkage and longer survival times than typically occur with other therapies [5, 6, 4, 7]. However, not every patient responds to treatment with ICIs. In fact, responders make up the minority of patients. Estimates of ICI response rates (the percentage of patients who meet some response criterion such as RECIST [8]) are in the range of 20-45% [7]. Additionally, like other targeted therapies, it is possible for tumors to become resistant to treatment after an initial period of response [9].

In order to improve the precision with which ICIs are prescribed, many researchers have focused on identifying biomarkers predictive of patient response to these therapies. Many candidate biomarkers have been identified across many “omics” modalities including bulk RNA-seq data [10, 11], single cell RNA-seq (scRNA-seq) [12], the tumor micro-environment [13, 14, 15], and somatic mutations [16]. Candidate biomarker identification is still an active area of research where so far little consensus has been reached. In fact, a recent large-scale meta-analysis revealed that the expression levels of the genes CXCL9 and CXCL13 and the non-synonymous tumor mutational burden (TMB) are the only consistently effective biomarkers [17].

Most biomarker discovery papers take the response phenotype as a given and attempt to find biological measurements that separate the population of responders from the population of nonresponders. In this paper we take a different approach. We essentially ignore the labeling of patient subpopulations as responders and non-responders and aim instead to infer a novel patient stratification in an unsupervised manner using discrete Ollivier-Ricci Flow (ORF) [18, 19, 20]. We evaluate this stratification by looking at the the pairwise log-rank test between the progression free survival (PFS) curves [21] of the resulting partition. In cases where the differences between the PFS curves were statistically significant (log-rank *p* ≤ 0.1) we performed extensive analysis of the resulting stratified patient subpopulations and identified potentially prognostic biomarker genes and gene modules.

The chief contributions of this paper are the clustering method deployed and the resulting biomarkers. In the community detection literature the use of ORF for partitioning known graphs (as opposed to our case where the graph is constructed) has theoretical guarantees and has demonstrated state-of-the-art performance on common benchmarks [18, 22]. Noting that for certain classes of graphs the discrete curvature converges to the continuous curvature [23], we endeavored to apply discrete Ricci curvature flow with surgery to split specific data points. Ours is the first use of the ORF algorithm to cluster patients based on RNA-seq measurements. In fact, our approach is, to the best of our knowledge, the first use of ORF to cluster data points in any setting. We also provide confirmatory results on ICI therapy response biomarkers that have only recently been reported in the literature.

## 2 Results

### 2.1 Patient Clustering with Discrete Ricci Flow

Our method, which is sketched in Figure 1, is grounded in the so-called “manifold-hypothesis” [24]. Specifically, we assume that high dimensional vectors that correspond to patient-derived gene expression measurements live on some unknown lower dimension surface (see Figure 1,top). We further suppose that, in general, patients who will respond similarly to ICI therapy will be closer together on this hypothetical surface. If the surface were known, we would still need a way to partition it. In the field of Riemannian geometry, the notion of ‘Ricci Flow with surgery’ is one such method that has been used to great success, most famously being a core tool in proving the Poincaré conjecture [25, 26, 27]. We base our method on the view that approximating the surface, and the Ricci flow process, should yield a good partition of the data points. Concretely if we approximate the surface in the form of an undirected weighted graph and the Ricci flow process in an iterative way, then we should be able to meaningfully stratify patients in an *unsupervised* manner (without explicit reference to clinical outcome data). We now sketch the method a little more formally. The full definitions are given in Section 4.

**Figure 1:**
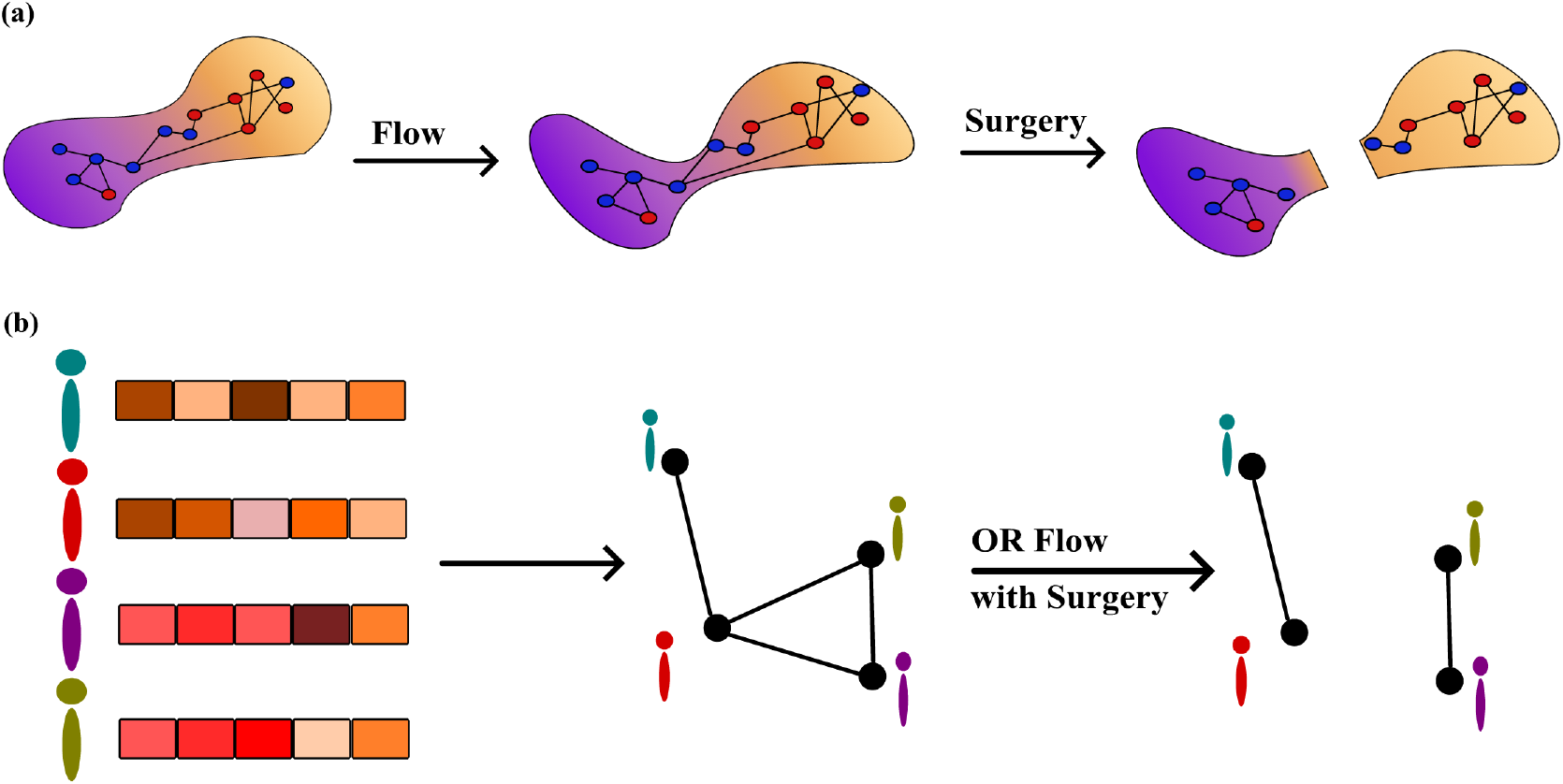
A schematic depicting the underlying point of view of our method **(a)** and the way in which it is implemented on a patient cohort **(b)**. Our assumption **(a)** is that patients’ transcriptomic measurements (blue and red dots) live on some unknown abstract surface (purple and orange blob). If responders (blue dots) and non-responders (red dots) are mostly in distinct regions of the surface then cutting the surface via Ricci flow should segregate the responder and non-responder populations. **(b)** In our method we approximate this unknown surface by first reducing the dimensionality of the transcriptomic measurements with locally linear embeddings (LLE) and then constructing a *k*-nearest neighbors graph where each vertex corresponds to a patient. We then partition the vertices/patients using discrete ORF with surgery.

We assume access to a *patient cohort*, which we take to mean a collection of patients with cancers from the same tissue of origin who receive the same ICI treatment. A cohort of *n* patients contains *n* vectors ***x***^(1)^, …, ***x***^(*n*)^, each with *D* dimensions (***x***^(*j*)^ ∈ ℝ^*D*^) where ***x***^(*j*)^ is the gene expression measurements for patient *j*. We also assume that each patient has clinical data available in the form of a binary measure of response (responder or non-responder) as well as time and censorship status for progression free survival (PFS).

As our method assumes an underlying surface we perform a manifold learning step to reduce the dimensionality of the data and approximate the assumed surface. Specifically we used locally linear embeddings (LLE) [28] to take the original collection of vectors ***x***^(1)^, …, ***x***^(*n*)^ to a new collection 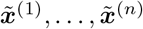 where 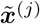 is the *d*-dimensional vector (*d* much smaller than *D*) corresponding to patient *j* after LLE.

Using the collection of vectors 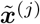 we construct a *k*-nearest neighbor graph where each vertex *j* corresponds to a specific patient. We assign each edge {*j, ℓ*} between patients/vertices *j* and *ℓ* in the graph a positive weight *w*(*j, ℓ*) and the discrete Ollivier-Ricci Flow (ORF) algorithm is then applied to partition the weighted graph.

The ORF algorithm proceeds roughly as follows. For a positive integer *T* we perform the following update *T* times:

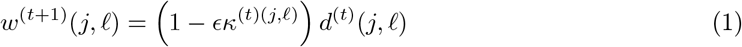

where *w*^(*t*)^(*j, ℓ*) is the weight assigned to edge {*j, ℓ*} at step *t, ϵ* > 0 is a step-size parameter, *κ*^(*t*)^(*j, ℓ*) is the Olliver-Ricci (OR) curvature of the edge at step *t*, and *d*^(*t*)^(*j, ℓ*) is the distance between vertices *j* and *ℓ* at step *t*. These terms are fully defined in Section 4. It is known that *κ* is negative on edges between tightly connected communities and positive within those communities [19, 29]. With this in mind the update in Equation (1) increases the weights of edges between communities and decreases the weights of edges within communities. After *T* steps we remove the edges with the largest weights and assign vertices/patients to groups corresponding to the remaining connected components.

### 2.2 Application to Two Patient Cohorts

We focused our analysis on two cohorts of patients treated with pembrolizumab (pembro). One cohort consisted of 45 patients with stomach adenocarcinoma (STAD) and the other was a cohort of 129 skin cutaneous melanoma (SKCM) patients. Additional experiments on patients with SKCM treated with nivolumab (nivo), another PD-1 targeted ICI, are presented in the Supplementary Material.

The results for partitioning the two pembro-treated cohorts are presented in Figure 2. Our method finds two subpopulations among both the STAD and SKCM cohorts which have meaningfully different survival profiles (log-rank p-values of 0.01 and 0.02, respectively). While the connected components shown in Figure 2 are mixed and generally reflect poor clustering from the perspective of binary response, our method did succeed in finding subpopulations with different survival profiles. The clustering displayed in Figure 2 suggests that the clustering of patients on the underlying surface is more reflective of the aggressiveness of the disease under treatment rather than response (i.e. tumor shrinkage in response to therapy).

**Figure 2:**
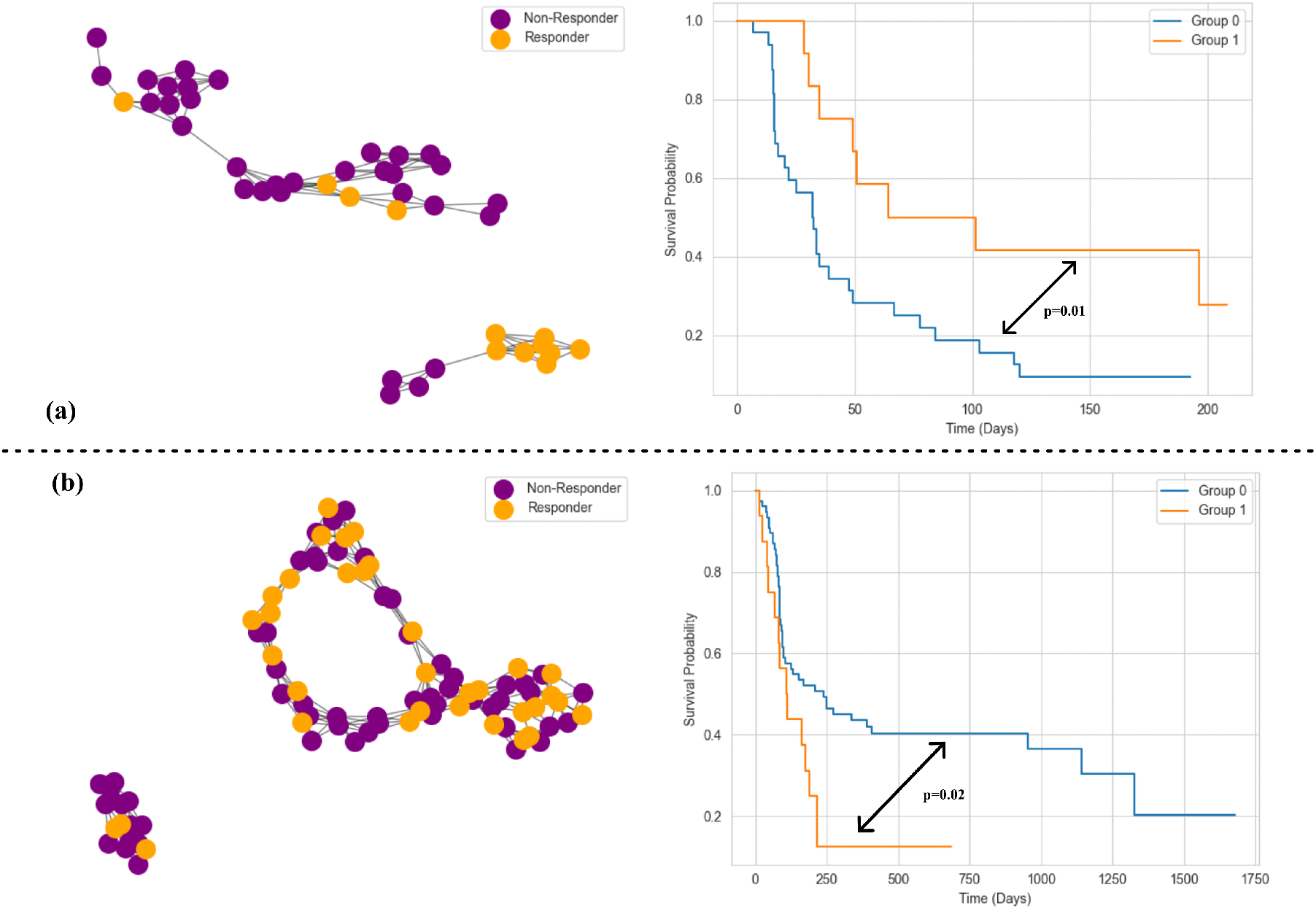
The results of partitioning two pembrolzumab treated cohorts of patients. The top row **(a)** gives the results in terms of the binary response labeled graph and the Kaplan-Meier fit survival curves for the STAD cohort. The bottom row **(b)** gives the same results for the SKCM cohort.

Next we investigated which genes could account for the distinct survival profiles. In order to find those genes we performed differential expression analysis across the discovered groups using DESeq2[30]. In order to summarize the genes that were found to be differentially expressed we performed a network pathway enrichment analysis, the results of which are in Supplementary Table 1. Interestingly we find that the DE genes between the STAD subpopulations yield a collection of pathways involved in antigen presentation. Antigen presentation is a key part of the adaptive immune system that allows cytotoxic T-cells to recognize tumor cells, among other crucial functions [31]. In addition the robustness of antigen processing has been proposed as a marker for ICI effectiveness [32, 33]. Indeed recent work has attempted to create an antigen presentation score for predicting ICI response in gastic cancer [34]. As antigen presentation is the means by which the immune system “discovers” tumor neoantigens, enriched antigen presentation pathways lend credence and nuance to the importance of neoantigen load as an ICI response biomarker [35, 33]. In the SKCM cohort we found a gene set enriched for immune response signaling, among others that are related to protein modification.

To assess if any of the discovered DE genes could be used as predictive response biomarkers we compared the distribution of expression among these DE genes between responders and non-responders. We used the Mann-Whitney U-test to compare the expression distributions among responders and non-responders with *p*-values adjusted for multiple hypothesis testing with the Bejamini-Hochberg procedure [36, 37] keeping those genes with an adjusted p value below 0.05. The expression distributions of these genes is displayed in Supplementary Figure 1.

Among the candidate biomarkers we found in this manner, many were known cancer-related genes and some were only recently discovered as being implicated in ICI response. For example *PLAAT4* is a known tumor supressor [38] but recent work implies it has a role in shaping the tumor-immune microenvironment in pancreatic cancer [39]. *MUC20* is another gene that can contribute to sculpting the micronenvironment. In fact in Supplementary Figure 1 the expression levels of *MUC20* are much higher among non-responders, which mirrors existing literature that shows that high *MUC20* expression levels are negatively correlated with positive clinical outcomes [40, 41, 42, 43].

In the interest of finding scalar-valued biomarkers of response, we partitioned the collection of DE genes into modules using hierarchical clustering. Among the DE genes discovered for STAD patients treated with pembrolizumab we found two modules (Figure 3). We performed leave-one-out cross validation using the module score (averaged expression across the module) as a single value predictor for response in a regularized logistic regression model. Both module 1 and module 2 displayed predictive value achieving accuracies of 68% and 66%, and area under the receiver operator curve (ROC/AUC) values of 0.73 and 0.7, respectively.

**Figure 3:**
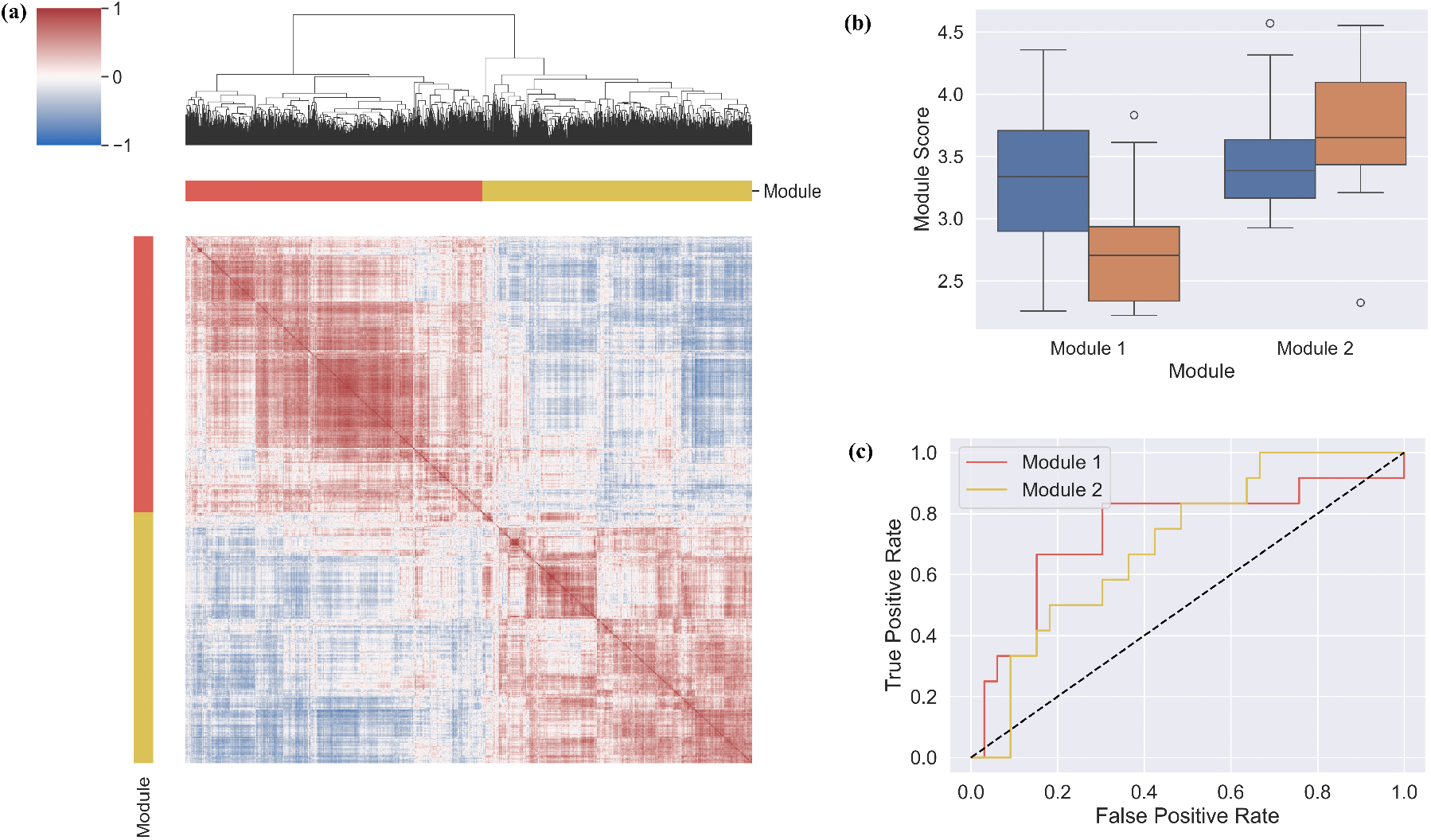
An analysis of the differentially expressed genes between the two discovered subpopulation of STAD patients treated with pembrolizumab. **(a)** A heatmap of the correlations between the gene expression values of the identified differentially expressed genes. The color bars show the modules found via hierarchical clustering performed using the displayed dendrogram above the heatmap. **(b)** Boxplot of the module scores (average expression in log_2_(transcripts per million+1) for responders and non-responders for each of the detected modules. In **(c)** we display the ROC curves for *ℓ*_2_-regularized logistic regression fit via leave-one-out cross validation (LOOCV). Module 1 had an area under the curve (AUC) of 0.74 while module 2 had an AUC of 0.7.

**Figure 4:**
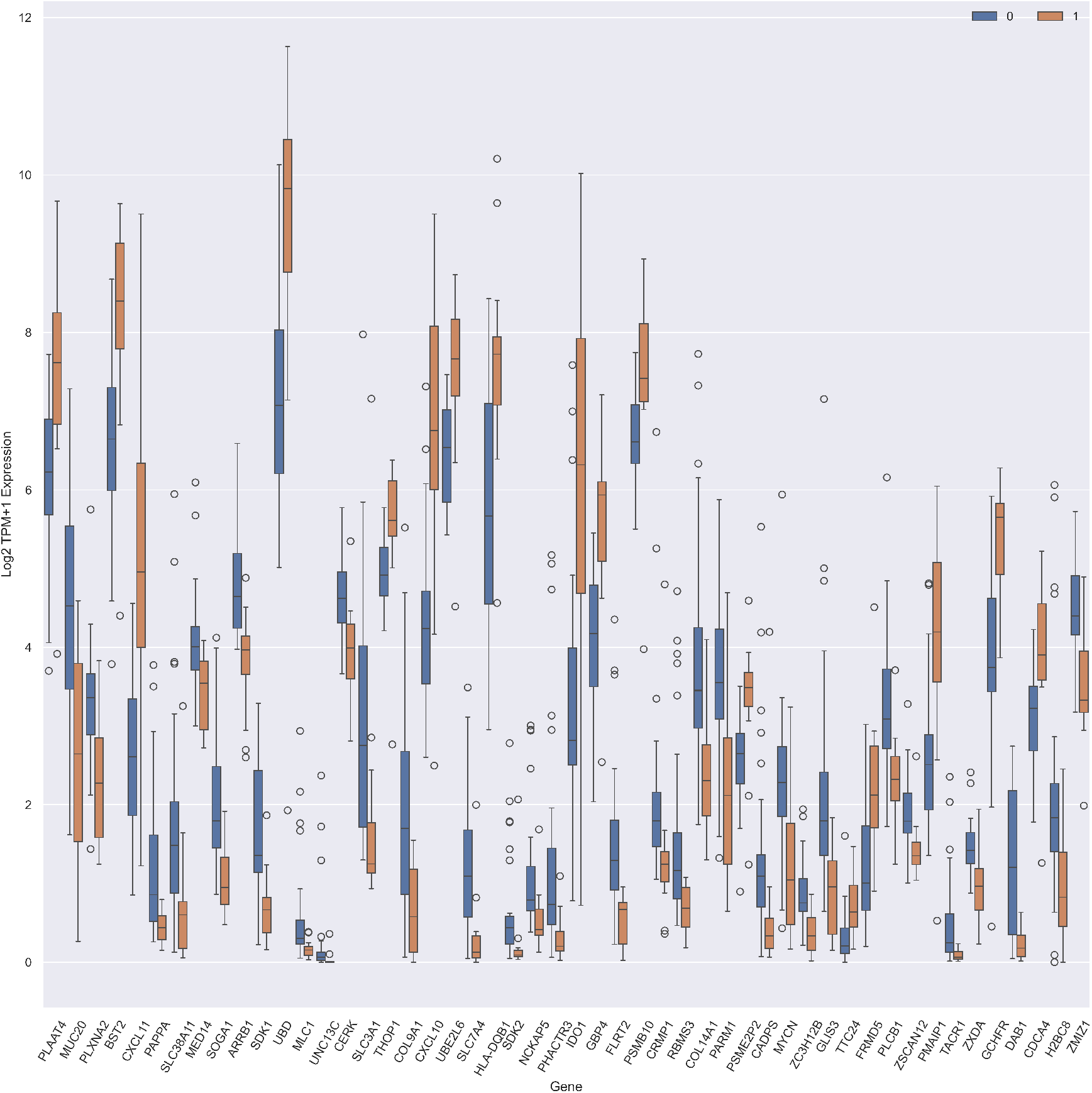
Genes found by performing differential expression analysis between our discovered groups in STAD patients treated with pembrolizumab. Displayed are only those genes which had statistically significant differences between expression distributions in responders and non-responders (multiple-testing adjusted Mann-Whitney p-value ≤ 0.05)

**Figure 5:**
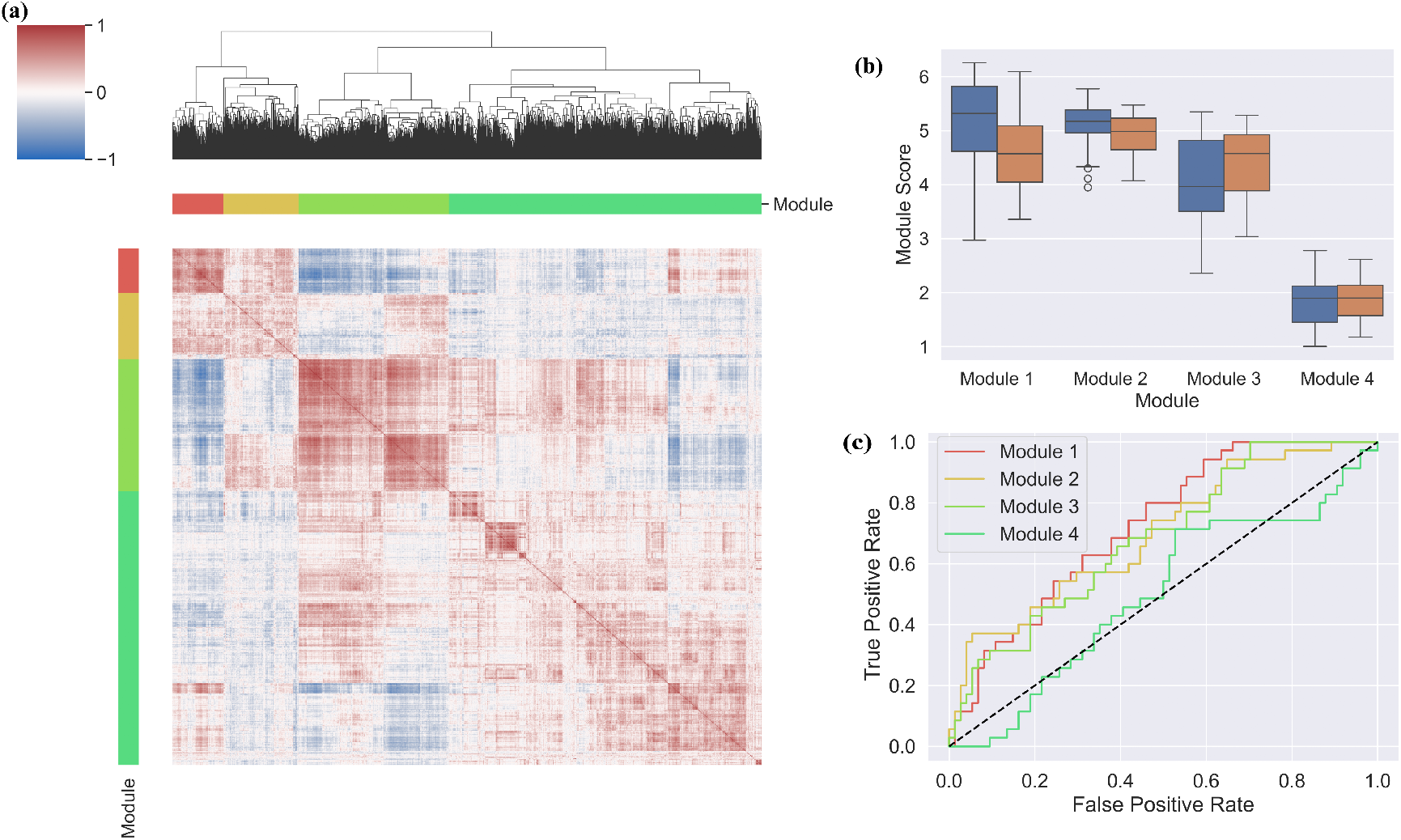
Results on module discovery for SKCM patients treated with nivolumab. **(a)** The the correlation heatmap for the expression of the genes found differentially expressed with the modules highlighted by the colored bars. **b** The module scores (average expression of genes in the module in log_2_(transcripts per million)+1) between responders and non-resonders. **(c)** The leave-one-out ROC/AUC for logistic regression using each module score as a single predictor. Module 1 had an ROC/AUC 0.72, and the values for Modules 2, 3, and 4 were 0.0.69, 0.68, 0.5, respectively

Similar experiments on the SKCM cohort yielded 3 modules, none of which demonstrated predictive value. In Supplementary Figure 2 and Supplementary Table 1, we present the results for the same methods applied to a cohort of SKCM patients treated with nivolumab. We found four modules with accuracies of 61%,63%, 59%, and 53% and ROC/AUC values of 0.72, 0.69, 0.68, and 0.5.

## 3 Discussion

We have presented a novel method for data clustering based on discrete Ollivier-Ricci flow. We found that approaching data clustering in the context of predicting patient response to lead to meaningful unsupervised discoveries of patient subpopulations. There are a number of potential directions for future work.

First we note that this method did not work for patients with kidney renal clear cell (KIRC) and bladder urothelial carcinoma (BLCA). An immediate next step would be to investigate if the best manifold learning method is drug or tissue dependent, or an artifact of the dataset. It would be valuable to check if other manifold learning techniques, such as t-SNE, UMAP, or Laplacian eigenmaps would be more effective. We also have ongoing work investigating whether the recently developed techniques for building *learnable* geometries [44] can lead to improved performance in ICI response prediction.

Additionally we hope to develop performance guarantees for our method. It is known that in a limited version of the stochastic block model, a generative model of graph formation, the ORF flow approach can perfectly recover communities [18]. One avenue of research that is worthy of exploring would be extending the work of [23], perhaps with reference to the *geometric* block model [45], to determine how the curvature behaves in a randomly generated nearest neighbor graphs. Relatedly, it is, to our knowledge, unknown if the ORF procedure truly reflects the structure of its continuous analogue. That is: do convergence results like those in [23] hold for discrete Olliver-Ricci flow.

We also could consider applying this method in a semi-supervised context. Namely, if only a subset of our data points have labels, the algorithm here could be used to cluster all the data points and assign labels to the unlabeled points by looking at the connected component in which it resides. On the bioinformatic side confirmatory studies of our candidate biomarkers, as well as the discovered modules, performed *in vitro* in mouse models or *in silico* using larger datasets would be useful for giving our work additional translational impact.

## 4 Methods

### Data Collection and Preprocessing

We downloaded raw data from the CRI-iAtlas [46] which is a publicly accessible repository of patient-derived bulk RNA-seq data and tumor-immune microenvironment measurements paired with clinical outcome data for humans treated with ICIs. We only used data from those samples taken **before** ICI treatment was administered and for which progression free survival (PFS) data (time until progression and censorship status) and clinical measures of response data were available.

Clinical response measurements in the iAtlas fall into four distinct categories: complete response, partial response, progressive disease and stable disease. We binned these into two groups: responders — made up of patients with complete or partial response — and non-responders — those with progressive or stable disease.

We curated a collection of genes thought to be related to cancer and ICI response. For cancerrelated genes we collected the genes listed in the KEGG database [47] under the heading “KEGG: Pathways in Cancer”, available at https://www.genome.jp/pathway/hsa05200, and the genes listed in the landmark paper on cancer genome landscapes [48]. For the ICI-related genes we used the genes involved in the computation of the IMPRES score [10]. We kept only those genes that were in our curated set and also had measurements in the CRI-iAtlas. Gene expression values were quantified in terms of transcripts per million (TPM), which were re-scaled via the logarithmic transformation log_2_(TPM + 1). These values were used to construct were the original data vectors ***x***^(*j*)^ in each cohort.

### Graph Theory

We consider only undirected weighted graphs. Specifically a graph *G*, written *G* = (*V, E*) consists of a finite set *V* of elements called *vertices* and *E* ⊂ *V* × *V* of edges. A graph is undirected if the edges “go both ways”, i.e: (*u, v*) ∈ *E* implies (*v, u*) ∈ *E*. We write edges as {*u, v*}. If there is an edge between two vertices we write this as *u* ∼ *v*.

The weight of an edge is a positive value which we write as *w*(*u, v*). The *degree* of a vertex *v*, written *d*_*v*_, is the sum of the weights that are incident to it, that is: *d*_*v*_ = ∑ _*u*:*u*∼*v*_ *w*(*v, u*). A *path P* (*u, v*) between two vertices *v, u* is a sequence of vertices *q*_1_, *q*_2_, …, *q*_*r*_ where no vertex appears more than once and *q*_1_ = *v, q*_*r*_ = *u* and *q*_*i*_ ∼ *q*_*i*+1_. The *length* of a path *P* (*u, v*), ℒ (*P* (*u, v*)) is taken to be the sum of the edge weights. Specifically:

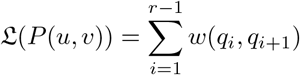

Lastly, for the distance between two vertices *u, v* we consider the all-pairs shortest path (APSP) distance. If 𝒫 (*u, v*) is the set of all paths between *u* and *v*, then we define the distance *d*(*u, v*) between *u, v* as

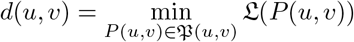

### Constructing the K-nearest Neighbors Graph

We have a collection of *n* patients represented as feature vectors ***x***^(1)^, …, ***x***^(*n*)^ where each ***x***^(*j*)^ has *D* dimensions representing expression levels for *D* distinct genes. We denote the expression level of gene *g* in patient *j* with 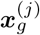. The LLE algorithm proceeds in two steps. The first step involves learning a set of weights Ω_*ij*_ between the vectors such that each ***x***^(*j*)^ can be approximately reconstructed as the sum of its neighbors. Specifically the weights are learned to minimize the function:

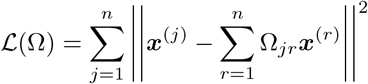

where || · ||^2^ represents the standard Euclidean distance between vectors, i.e:

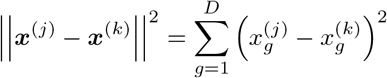

The set of weights Ω is found by solving an optimization problem subject to some constraints. Specifically one must specify the number *L* of neighbors to be considered in reconstructing each ***x***^(*j*)^ and then Ω is forced to satisfy: Ω_*ji*_ = 0 if ***x***^(*i*)^ is not one of ***x***^(*j*)^’s *L* nearest neighbors and that ∑_*j*=1_ Ω_*ij*_ = 1. We fixed *L* = 5 in our experiments.

The second step of LLE involves choosing an embedding dimension *d* < *D* and constructing reduced vectors 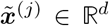 using the learned weights Ω. The embedding is found such that the *reconstruction error*

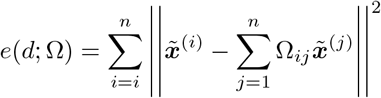

is minimized. Details on this procedure can be found in [49]. In our experiments we tried every *d* in the range 2, …, 15 and used the one for which the reconstruction error was the smallest, which lead to *d* = 2 for all cohorts.

To construct the patient-patient graph we apply a symmetrized *K*-nearest neighbors. Let *N*_*K*_(*j*) be collection of indices *i* ∈ {1, …, *n*} for which 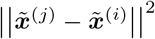 is the smallest. We then add a vertex for each patient and an edge {*j, ℓ*} between patients *j* and *ℓ* according to the rule

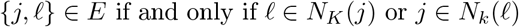

The initial weights of the edges were all set to *w*(*j, ℓ*) = 1 as setting the weights to distances made no difference in our experiments.

### Discrete Ollivier-Ricci Flow with Surgery

The notion of graph curvature that we use was developed in a landmark paper by Ollivier [50] in the more general setting of Markov chains on metric spaces. In this work we adopt the form of this curvature, known as Ollivier’s Ricci Curvature (ORF), developed by Lin and Yau [51]. We now describe how to construct this curvature and perform graph partitioning with ORF flow on a general undirected weighted g raph. The s teps are performed identically on the specific patient-patient interaction graph.

We begin with a value *α* ∈ [0, 1] and assign a one-step random walk probability to each vertex *v*. For any given *α* and any vertex *v* the probability distribution 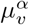 is given by

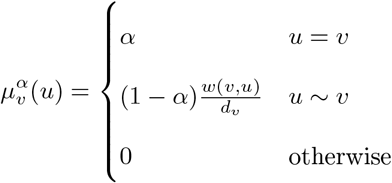

The curvature *κ*(*u, v*; *α*), for any given *α* and edge *u, v*, is compute via:

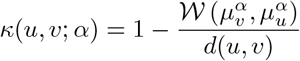

where 𝒲 is the well-known Wasserstein distance [52, 53]. Since in all our experiments *α* = 0 we now drop *α* from the notation and write the curvature as *κ*(*u, v*). Let *d*_max_ be the largest **number** of neighbors that any vertex in the graph has. The computational complexity of computing *κ*(*u, v*)) is 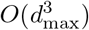 as *d*(*u, v*) can be pre-computed in *O*(|*V* |^3^) time and 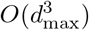 is the computational complexity of computing the Wasserstein distance via a linear programming.

The base of our method, Algorithm 1, is the discrete Ollivier-Ricci Flow (ORF) procedure [19, 20, 18] which is based on the update given in Equation (1). At each iteration we must recompute the pairwise distances and curvature values, which for *T* iterations has complexity 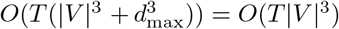. For all drugs and tissues we set *ϵ* = 0.3, *K* = 5, *T* = 50.

The ‘surgery’ portion of the algorithm involves cutting the weights that remain after the *T* iterations. We use the heuristic procedure of [22]. Let be *w*_max_ be the largest edge weight in the graph and *w*_min_ a smallest weight value which is left as a parameter and *δ* > 0 a step size. We iterate over a grid of values *w* between *w*_max_ and *w*_min_ equally spaced by *δ* in decreasing order, removing edges with weight greater than *w*. At each step we compute the modularity *Q*(*w*) [54] of assigning each node to a community that corresponds to the connected component in which it lies.

### Differential Expression and Pathway Enrichment

For differential expression we used the DESeq method from the R package of the same name [30]. We kept all genes which were differentially expressed with an adjusted Wald p-value below 0.01 as output by the DESeq2 software. For network-based enrichment we used the WebGestalt tool [55] to perform network-based pathway analysis. The “method of interest,” “organism of interest,” and “functional database” parameters were set to “Network Topology-based Analysis”, “homo sapiens”, and “network”/”PPI BIOGRID” respectively.

### Implementation

Code to perform these experiments was written in python. We used the library scikit-learn [56] to perform LLE and fit the logistic regression models. Code supporting this can be found at https://github.com/jbannon/ORFlowICIs.

#### Algorithm 1

Data Clustering with Ollivier Ricci Flow

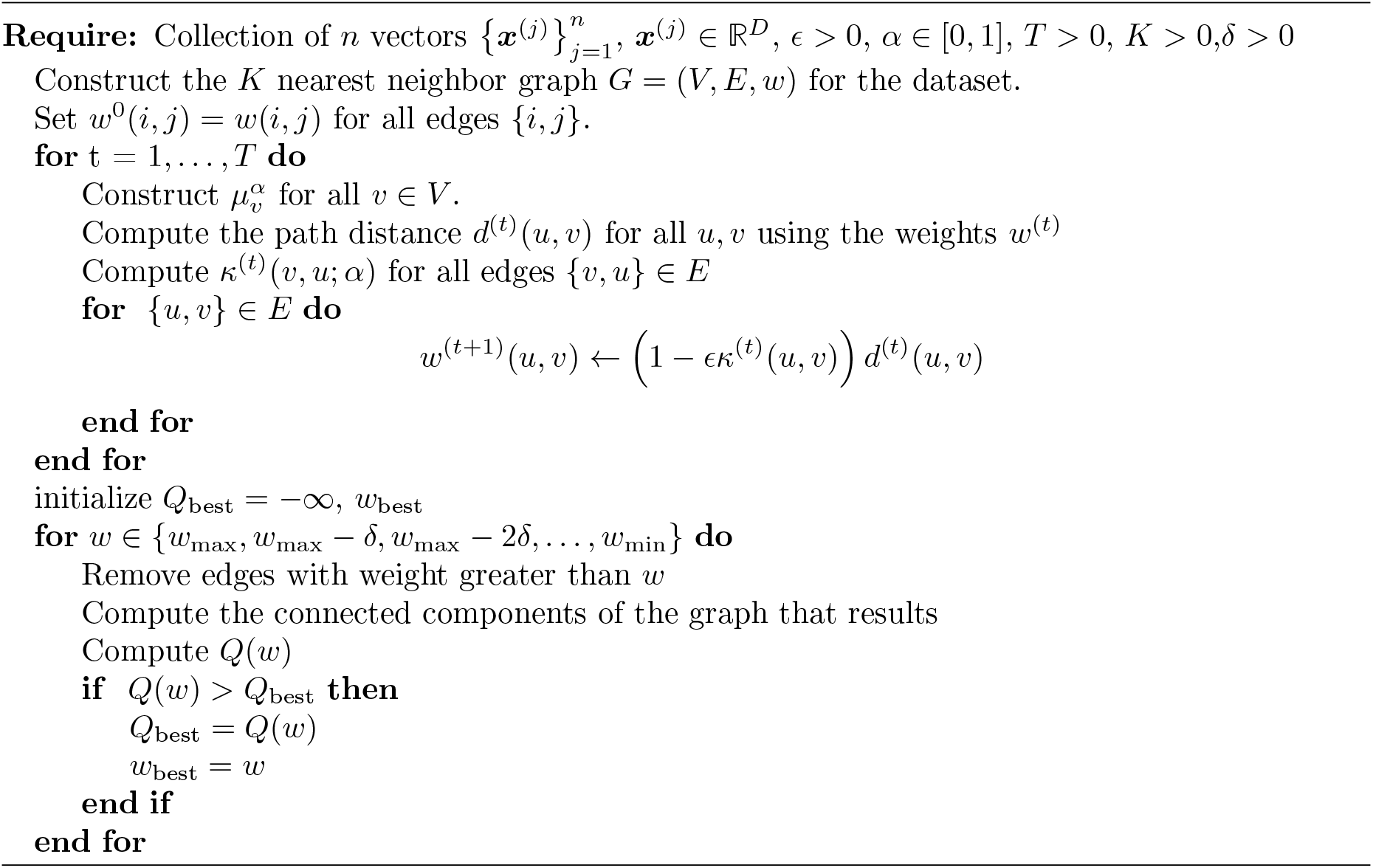

## 5 Data Availability Statement

The data underpinning our experiments is publicly available through the CRI-iAtlas. Specifically one can navigate to the portal at https://cri-iatlas.org and download the data from there. The supporting code is at https://github.com/jbannon/ORFlowICIs.

## 6 Acknowledgements

JB and BM are grateful to NYU’s departmental support during the early stages of this manuscript. JB is also grateful to many fruitful discussions with Dr. Andrew Hall.

## 7 Author Contributions

JB and CC conceived of the experiments. JB implemented the code and performed the analysis and figure generation and wrote the majority of the manuscript. JB, CC, and BM all edited the manuscript. All authors have read and approved the manuscript.

## A Supplementary Material

### Additional Results for Pembrolizumab

**Table 1:** Network-based pathway enrichment analysis for pembrolizumab patients. Enriched pathways for patients with STAD are the left and those for SKCM patients are on the right.

| Pembro - STAD |  | Pembro - SKCM |  |
| --- | --- | --- | --- |
| GO ID | Description | GO ID | Description |
| GO:0002478 | antigen processing and presentation of exogenous peptide antigen | GO:0043170 | macromolecule metabolic process |
| GO:0048002 | antigen processing and presentation of peptide antigen | GO:0036211 | protein modification process |
| GO:0019886 | antigen processing and presentation of exogenous peptide antigen via MHC class II | GO:0006807 | nitrogen compound metabolic process |
| GO:0002495 | antigen processing and presentation of peptide antigen via MHC class II | GO:0002764 | immune response-regulating signaling pathway |
| GO:0002399 | MHC class II protein complex assembly | GO:0002757 | immune response-activating signaling pathway |
| GO:0002503 | peptide antigen assembly with MHC class II protein complex | GO:0002253 | activation of immune response |
| GO:0002396 | MHC protein complex assembly | GO:0043412 | macromolecule modification |
| GO:0002501 | peptide antigen assembly with MHC protein complex | GO:0090304 | nucleic acid metabolic process |
| GO:0019884 | antigen processing and presentation of exogenous antigen | GO:0044238 | primary metabolic process |
| GO:0002504 | antigen processing and presentation of peptide or polysaccharide antigen via MHC class II | GO:0071704 | organic substance metabolic process |

### Results for SKCM Patients Treated with Nivolumab

**Table 2:** Gene ontology (GO) pathway enrichment for SKCM patients treated with nivolumab.

| GO ID | Description |
| --- | --- |
| GO:0048583 | regulation of response to stimulus |
| GO:0034097 | response to cytokine |
| GO:0071345 | cellular response to cytokine stimulus |
| GO:0006915 | apoptotic process |
| GO:0071310 | cellular response to organic substance |
| GO:0019538 | protein metabolic process |
| GO:0070887 | cellular response to chemical stimulus |
| GO:0008219 | cell death |
| GO:0006412 | translation |
| GO:0006950 | response to stress |

